## Supplementary material for "Interphase chromosomes of the *Aedes aegypti* mosquito are liquid crystalline and can sense mechanical cues": SI

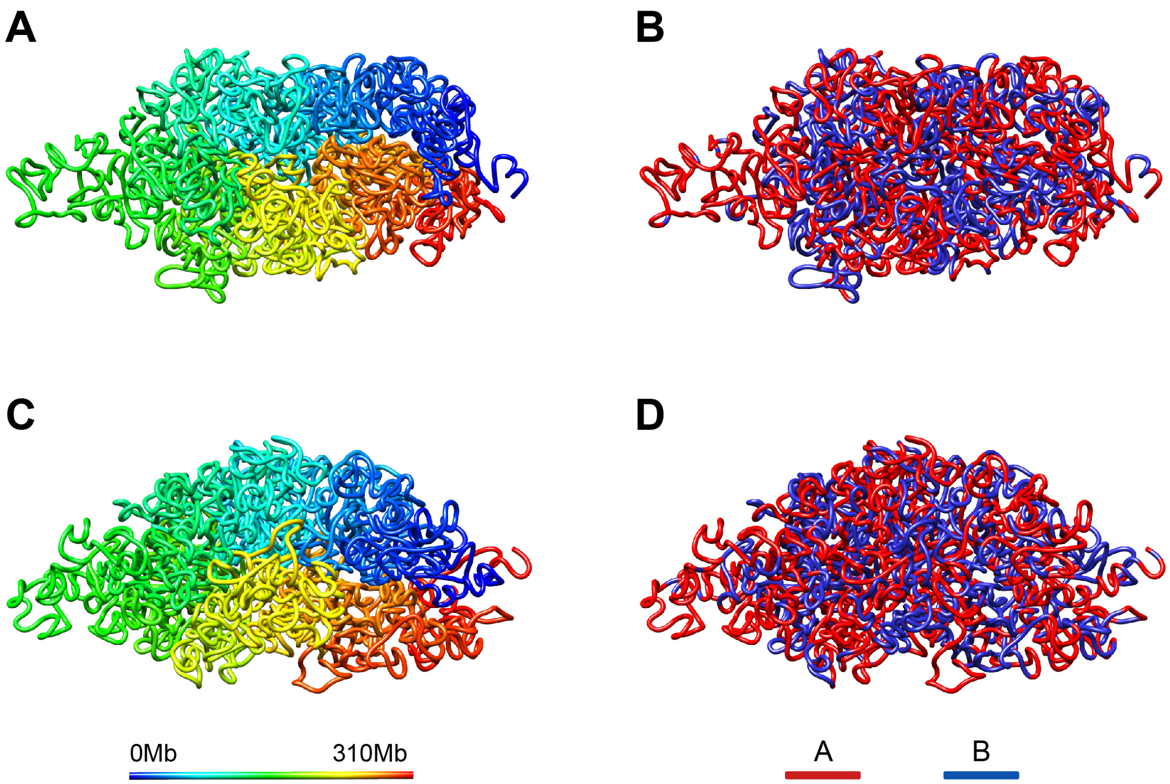


Figure S1: Representative 3D structures of chromosome 1 from *Aedes aegypti* modeled at the nominal information theoretic temperature ($T = 1.0$)[1]

.

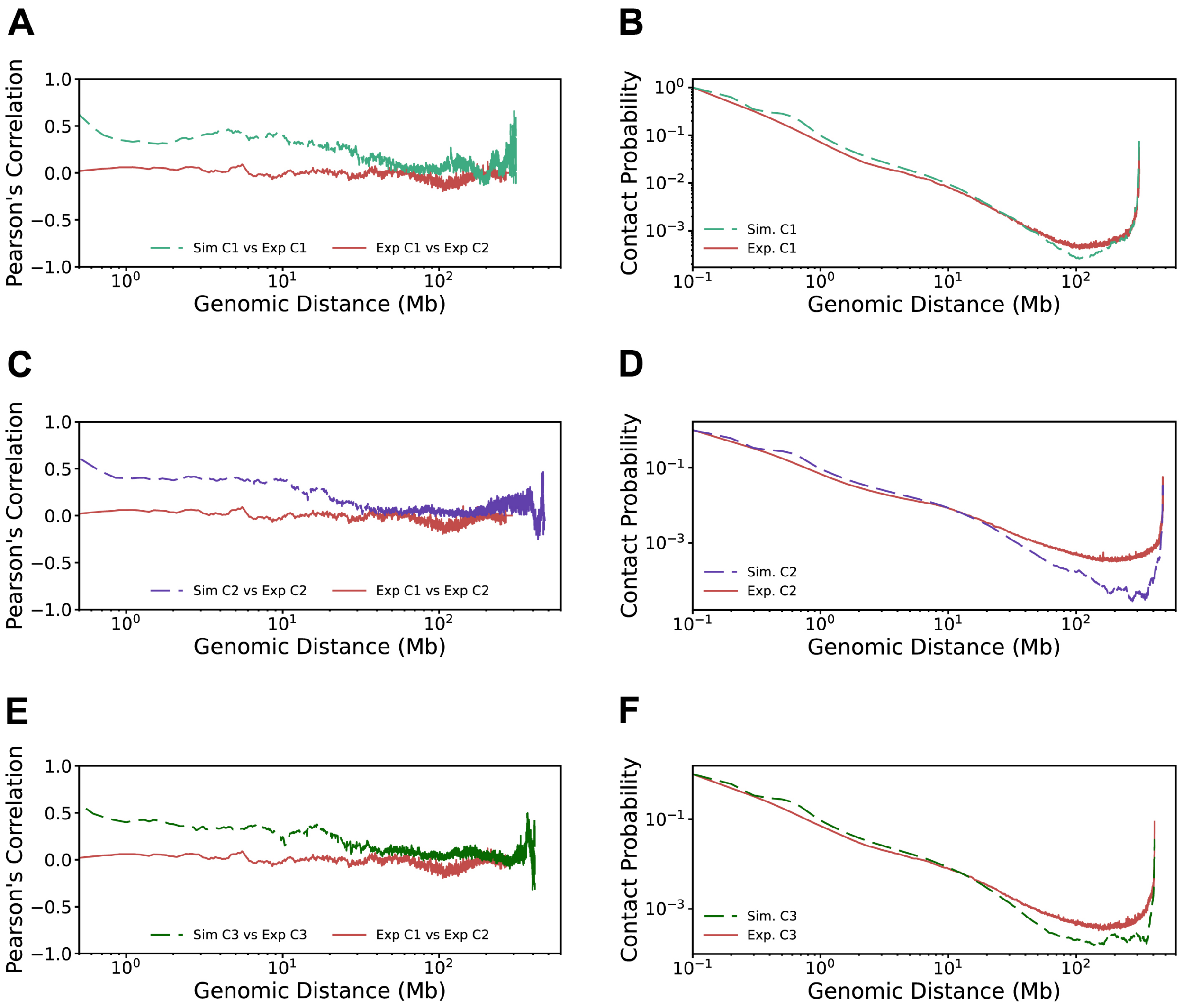


Figure S2: Pearson's correlation and contact probability as a function of the genomic distance.[2–5] The curves were obtained for each chromosome extracted from the *Aedes aegypti* full nucleus simulation.

Table S1: Kolmogorov-Smirnov statistic two-sided test (KS-value) comparing the High and Low ATAC-seq value distributions. The test computes the null hypothesis that two independent data set are obtained from the same continuous distribution.[6] The distributions are presented in Figure 4D of the main text.

| **Chromosomes** | **KS-value** |
|---|---|
| 1P and 1M | 0.26 |
| 2P and 2M | 0.52 |
| 3P and 3M | 0.27 |
| Whole Genome | 0.58 |

### Method Details

#### Homopolymer model

The homopolymer potential models a generic bead-spring polymer in which each bead represents a genomic segment of 100 Kb in sequence. The potential energy $U_{HP}(\vec{r})$ describes a spatially self-avoiding polymer (no nonbonding attractive term is included here) and serves as a support for the features added by using the maximum entropy principle. This potential consists of the following five terms, $U_{\text{FENE}}$ , $U_{\text{Angle}}$ , $U_{hc}$ and, $U_{sc}$:

$$\begin{aligned} U_{HP}(\vec{r}) &= \sum_{i\in\{\text{Loci}\}} U_{\text{FENE}}(\vec{r}_{i,i+1}) + \sum_{i\in\{\text{Loci}\}} U_{hc}(\vec{r}_{i,i+1}) + \sum_{i\in\{\text{Angles}\}} U_{\text{Angle}}(\theta_i) \\ &+ \sum_{\substack{i,j\in\{\text{Loci}\}\\ j>i+2}} U_{sc}(\vec{r}_{i,j}), \end{aligned} \tag{1}$$

where,

$$U_{\text{FENE}}(\vec{r}_{i,j}) = \begin{cases} -\frac{1}{2}k_b R_0^2 \ln\left[1 - \left(\frac{r_{i,j}}{R_0}\right)\right] & r_{i,j} \leq R_0 \\ 0 & r_{i,j} > R_0 \end{cases}$$

$U_{\text{FENE}}$ (Finite Extensible Nonlinear Elastic potential) is the bonding term applied between two consecutive monomers.

$$U_{hc}(\vec{r}_{i,j}) = \begin{cases} 4\epsilon\left[\left(\frac{\sigma}{r_{i,j}}\right)^{12} - \left(\frac{\sigma}{r_{i,j}}\right)^{6} + \frac{1}{4}\right] & r_{i,j} \leq \sigma 2^{\frac{1}{6}} \\ 0 & r_{i,j} > \sigma 2^{\frac{1}{6}} \end{cases}$$

$U_{hc}(\vec{r}_{i,j})$ is the hard-core repulsive potential, include to avoid overlap between bonded monomers.

$$U_{\text{Angle}}(\theta_i) = k_a[1 - \cos\Theta_i - \theta_0],$$

a three-body potential applied to all connected three consecutive monomers, where $\theta_i$ is the angle defined by two vectors $\vec{r}_{i,i+1}$ and $\vec{r}_{i,i-1}$.

The non-bonded pairs is defined by a soft-core repulsive interaction in the following form:

$$U_{sc}(\vec{r}_{i,j}) = \begin{cases} \frac{1}{2}E_{\text{cut}}\left[1+\tanh\left(\frac{2U_{LJ}(\vec{r}_{i,j})}{E_{\text{cut}}}-1\right)\right] & r_{i,j} \le r_0 \\ U_{LJ}(\vec{r}_{i,j}) & r_0 \le r_{i,j} \le \sigma 2^{\frac{1}{6}} \\ 0 & r_{i,j} > \sigma 2^{\frac{1}{6}} \end{cases}$$

The expression $U_{LJ}$ correspond to the Lennard-Jones potential $U_{LJ}(\vec{r}_{i,j}) = 4\epsilon\left[\left(\frac{\sigma}{r_{i,j}}\right)^{12} - \left(\frac{\sigma}{r_{i,j}}\right)^{6} + \frac{1}{4}\right]$ capped off at a finite distance, thus allowing for chain crossing at finite energetic cost. $r_0$ is chosen as the distance at which $U_{LJ}(\vec{r}_{i,j}) = \frac{1}{2}E_{cut}$

Centromeric and telomeric regions are anchored to the nucleus wall using a flat-bottomed potential $U_{\text{rfb}}$. This potential is employed to restrain the loci within a simulation volume. If a locus moves outside the chosen region, a harmonic force moves the bead to the flat-bottomed region. On the other hand, there is no force acting on the centromeric and telomeric locus within the flat-bottomed part of the potential. The flat-bottomed potential is described as follow:

$$U_{\text{rfb}}(\vec{r}_i) = -\frac{1}{2}k_{rfb}(r_i - R_0)^2\Theta(r_i - R_0) \tag{2}$$

where $R_0$ is the center flat bottom potential location, $r_i$ is the locus's position, $k_{rfb}$ the force constant, and $\Theta$ is the Heaviside step function.

#### Crosslinking Probability Function

The function $f(r_{i,j})$ is the probability of crosslink[1,2] and can be written as:

$$f(r_{i,j}) = \frac{1}{2}\left(1 + tanh\left[\mu(r_c - r_{i,j}\right]\right) \tag{3}$$

where $\mu$ ad $r_c$ are determined based on the experimental Hi-C maps. The function $f(r_{i,j})$ must return 1 when two beads are in contact (distance between the center of two beads is equal to 1, in reduced units $\sigma$), e.i., $f(1) = 1$. $f(r_{i,j})$ also must decreases monotonically with the distance and the minimum of the experimental probabilities must match with the next nearest neighbor, e.g, $f(2) = \min\{P^{exp}_{i,i+2}\}$. The parameters adjusted for the Hi-C maps of *Aedes aegypti* are $\mu = 3.48$ and $r_c = 1.76$.

Table S2: Type-to-type interaction values for the *Aedes aegypti* genome in units of KT.

| | A | B |
|---|---|---|
| A | -0.252 | -0.273 |
| B | -0.273 | -0.339 |
